# Preparation and in vitro release of fluorouracil polylactide glycolide-polyethylene glycol monomethyl ether nanoparticles

**DOI:** 10.1101/628792

**Authors:** Anil Shumroni, David Gupta

**Affiliations:** Department of Chemistry, University of Alabama

**Author notes:** Address: 250 Hackberry Ln, Tuscaloosa, AL 35401.

## Abstract

This report demonstrates a novel strategy to prepare fluorouracil polylactide glycolide-polyethylene glycol monomethyl ether (PLGA-mPEG) nanoparticles and study their in vitro release characteristics. Fluorouracil PLGA-mPEG nanoparticles were prepared by nanoprecipitation method. The encapsulation efficiency was determined by high performance liquid chromatography. Based on the single factor experiment, the prescription and preparation process were optimized by orthogonal experiments. The in vitro release characteristics of nanoparticles were studied by dynamic membrane dialysis. Results The prepared nanoparticles were relatively uniform spheroidal particles with an average particle size of about 124. 3 nm, a Zeta potential of - 20. 6 mV, and an average encapsulation efficiency of (44.72 ± 0.38%). In vitro drug release experiments showed that the particle burst release was less than 30% at 2 h, and the drug was slowly released within 48 h after burst release.

## Introduction

5-fluorouracil (5-Fu) is an anti-metabolic anti-tumor drug with a strong killing effect on proliferating cells. It is a broad-spectrum anti-tumor drug with a wide range of clinical applications. It is clinically used for the treatment of various tumors and has obvious curative effects on various cancers such as digestive tract cancer and ovarian cancer, cervical cancer and bladder cancer [1]. Most of the current clinical use is 5-Fu common injection. Because 5-Fu injection has low selectivity to tumor cells and is widely distributed in the body, it is more toxic to normal cells and is prone to many adverse reactions such as bone marrow. Inhibition, leukopenia, thrombocytopenia, etc. [2]. Therefore, in order to improve the targeting of 5-Fu, reduce adverse reactions, and prolong the duration of drug action, research and development of 5-Fu new drug delivery system has become a research hotspot in recent years.

Nanoparticles are a new type of drug delivery system developed in recent years, and their application in antitumor drugs has received extensive attention. Because the tumor tissue is rich in blood vessels, the nano-sized particles easily penetrate the blood vessels of the tumor site and remain in it, which has high permeability and retention (EPR), so the nanoparticles themselves have certain tumor targeting properties. At the same time, the nanoparticles also have the advantages of improving drug stability, prolonging the action time of the drug, and improving the bioavailability of the drug [3-4]. Polylactide-glycolide-polyethylene glycol monomethyl ether (PLGA-mPEG) has good biocompatibility and biodegradability as a novel carrier material [5-6]. The preparation of the drug cisplatin into PLGA-mPEG nanoparticles can reduce the toxicity of the drug, enhance the passive targeting of the drug, prolong the circulation time of the drug in the body, and thereby significantly improve the therapeutic effect of cisplatin [7]. In this experiment, PL-p-PEG was used to prepare 5-Fu nanoparticles by nanoprecipitation method, and its quality and drug release characteristics were studied to lay a new high-efficiency, low-toxic and long-acting new 5-Fu nanoparticle preparation. Experimental basis.

## Materials and Method

AEU - 210 Precision Electronic Balance (Shimadzu Corporation, Japan); CL - 2 Magnetic Stirrer (Gongyi Yuhua Instrument Co., Ltd.); RE - 52C Rotary Evaporator (Gongyi Yingying Yuhua Instrument Factory); SHB - D Type micro circulating water vacuum pump (Zhengzhou Changcheng Branch Industry and Trade Co., Ltd.); TGL-16 high speed desktop centrifuge (Shanghai Medical Instrument Factory); High Performance Liquid Chromatograph (Waters, USA); KQ3200E Medical Ultrasonic Cleaner (Kunshan Ultrasound) Instrument Co., Ltd.; Zetasizer 3000HS Laser Particle Size Analyzer (Malvern, UK); Tecnai G2 Transmission Electron Microscope (Philips, The Netherlands).

PLGA-mPEG (relative molecular mass 1 200, Shandong Jinan Biotechnology Co., Ltd.); 5-Fu (content ≥ 99. 5%) (Shanghai Senyu Fine Chemical Co., Ltd., batch number: 20081030); Polosham 188, poloxamer 407 (BASF, Germany); methanol is chromatographically pure; acetone, ethanol and other reagents are of analytical grade; distilled water (made by the Pharmacy Department of Harbin Medical University).

## Results and discussion

Single factor experiment The effects of the type of organic solvent, the temperature of the aqueous phase, the type and concentration of the surfactant, the amount of the polymer, the quality of the organic phase, and the stirring time on the encapsulation efficiency were investigated by single factor experiments.

On the basis of single factor experiments, four factors, ie organic phase composition, water bath temperature, drug loading ratio and surfactant concentration, which significantly affect the encapsulation efficiency of the drug, were optimized by orthogonal design. Three factors were selected for each factor, and experiments were performed according to L9 (34), and the best prescription was determined by the encapsulation rate as the main evaluation index. Orthogonal experimental design

**Figure 1.**
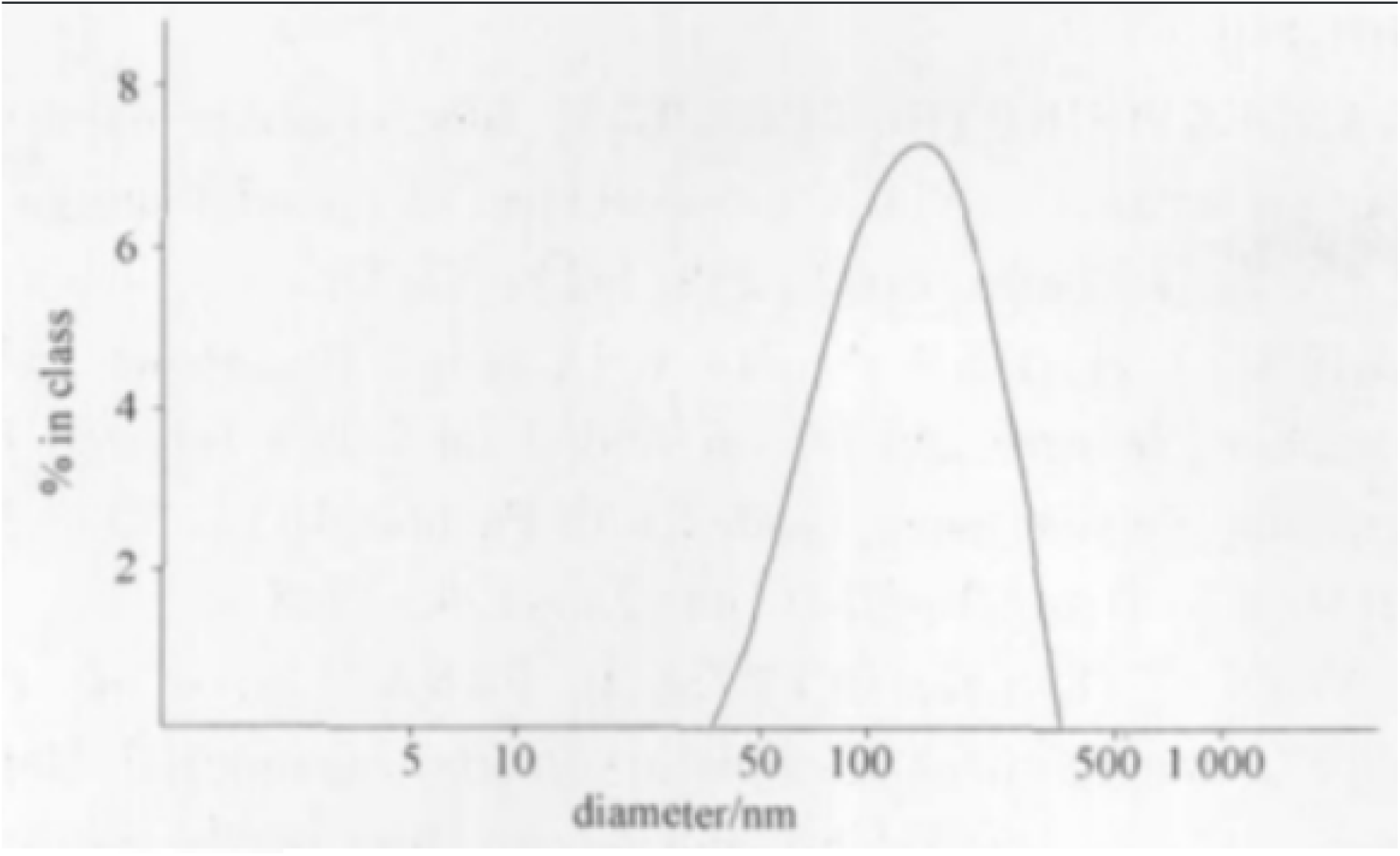
DLS analysis of nanoparticles.

It can be seen from Table 2 that this experimental calibration model P < 0.05 has statistical significance. Based on the results of Tables 1 and 2, the primary and secondary order of each index factor is A > B > C > D, and the two factors A and B have statistical significance (P < 0.05). In the factor A, kI > kII > kIII, B factor in kII > kIII > kI, C factor > kII > kI > kIII, and D factor kII > kIII > kI, so the best place is A1B2C2D2.

In this experiment, PLGA-mPEG was used as the carrier material to prepare nanoparticles by nanoprecipitation. 20 mg of 5-Fu and 50 mg of PLGA-mPEG were dissolved in 6 mL of organic solvent acetone at room temperature as an organic phase, and the organic phase was added dropwise at a constant rate to 40 mL of a 1% poloxamer stirred at 40 °C. The microporous membrane of μm was filtered, and the filtrate was centrifuged for 40 min (20 000 r min −1, 4 °C) to collect the nanoparticles to obtain 5-Fu nanoparticles.

Chromatographic conditions: Diamonsil TM C18 (4.6 mm × 250 mm, 5 μm); mobile phase: methanol-water (1:1); flow rate: 1. 0 mL min −1; column temperature: 30 °C; Detection wavelength: 265 nm; Injection volume: 10 μL. Under this condition, the retention time of 5-Fu is about 4. 8 min.

Weigh accurately 5-Fu about 10 mg in a 100 mL volumetric flask, dissolve it in the mobile phase, dilute to the mark, and mix to prepare a stock solution of 100 μg mL −1. Separately absorb the stock solution 1. 0, 3. 0, 4. 0, 5. 0, 6. 0 and 8. 0 mL, place in a 10 mL volumetric flask, dilute to the mark with mobile phase, and obtain a series of certain concentration. Standard solution. Taking the peak area (A) as the ordinate and linear regression with the mass concentration ρ (μg mL - 1) as the abscissa, the regression equation is obtained: A = 3. 557 0 × 104 ρ + 1. 941 5 × 104 (r = 0. 999 5). 5-Fu has a good linear relationship between 10 and 100 μg mL −1.

**Figure 2.**
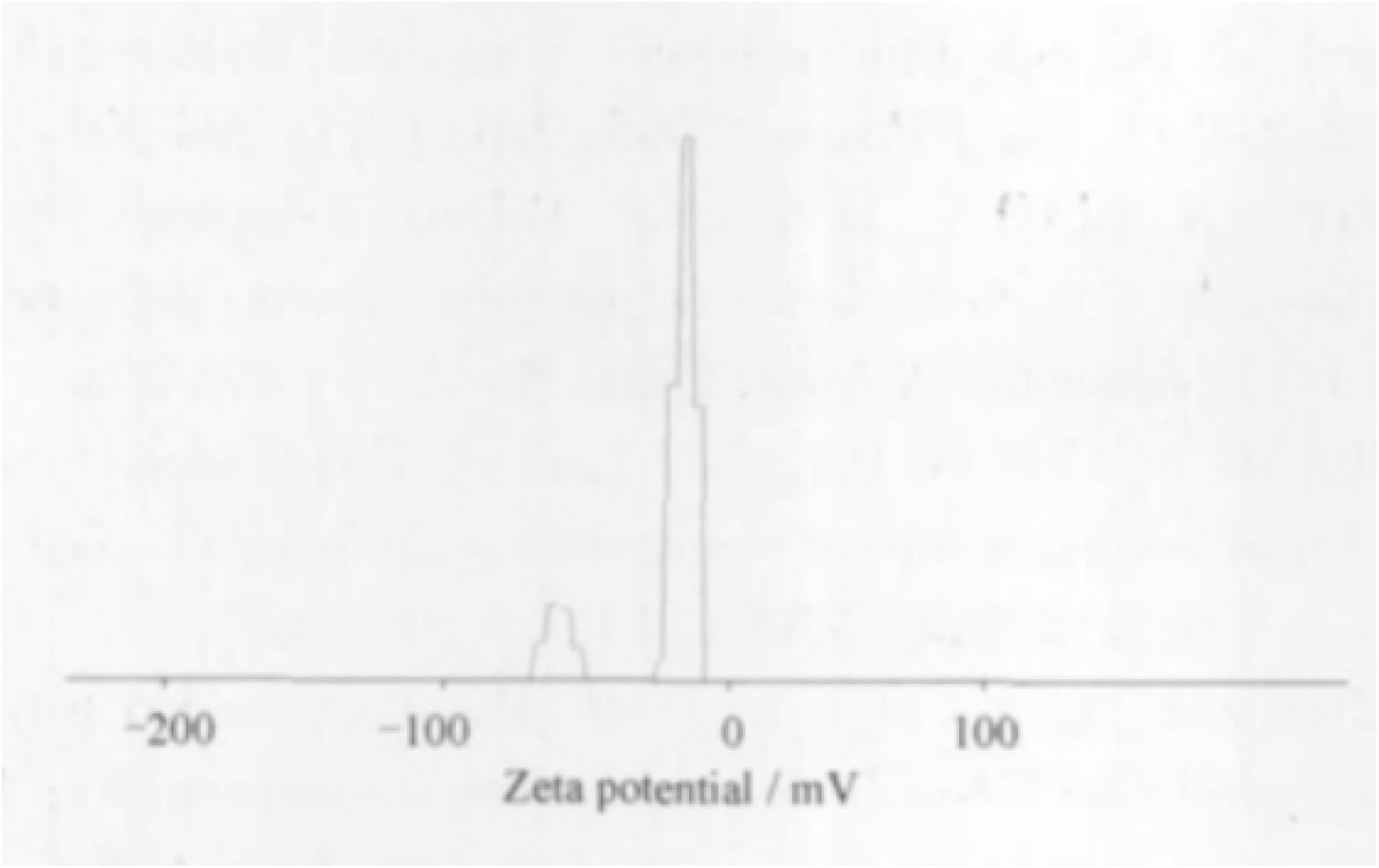
Zeta potential.

Take 40, 50, 60 μg · mL −1 low, medium and high concentration solutions of the reference substance, measure 5 times in 1 day, calculate intraday precision; inject once every 1 d, continuously measure 5 d, calculate the daytime precision. Intraday precision and daytime precision (RSD) are less than 2%, which meets the requirements.

Weigh 9 parts of a certain amount of blank nanoparticle preparation, add to the low, medium and high concentration solutions of 40, 50, 60 μg mL −1, and prepare 3 parts for each concentration. After it is completely dispersed, the supernatant is centrifuged for injection measurement, and the recovery rate is calculated. The average recovery was 100. 09%, and the RSD was 0.94%.

Determination of encapsulation efficiency The free drug and nanoparticles were separated by high-speed centrifugation, and the free drug content was determined by high performance liquid chromatography. After centrifugation of the drug-loaded nanoparticle solution for 40 min (20 000 r · min −1, 4 °C), the supernatant was collected, 10 μL was injected, the peak area was recorded, and the encapsulation efficiency was calculated.

Currently, biodegradable carrier materials commonly used for nanoparticles are polycyanoacrylates, polyesters, polyamides, polyanhydrides, etc. [8], of which polylactic acid (PL), polyglycolic acid and lactic acid/hydroxyl are the most widely used. Acetic acid copolymer (PLGA). It has been reported that PLA and PLGA can be used as carriers for drug-loaded microspheres and nanoparticles to protect drugs, solubilize, improve bioavailability and target drug release [9]. However, these two materials have severe hydrophobicity, which limits their use in the field of drug carriers. In this experiment, PLGA-mPEG was used as the carrier material. After the PLGA was attached to the hydrophilic segment mPEG, the hydrophilicity of the material was increased to make the material amphiphilic. At the same time, due to the introduction of hydrophilic PEG chain, it can resist and escape the phagocytosis of the reticuloendothelial system, delay the retention time of the circulatory system in the body, and facilitate the penetration of the drug into the vascularized tumor tissue, further improving the nanoparticle. Targeting. In addition, the nanoparticles prepared in this experiment have a certain sustained release property, which effectively reduces the large amount of release of the drug in the nanoparticles before being concentrated in the tumor tissue, thereby avoiding the systemic toxic side effects and the reduction of the drug effect.

**Figure 3.**
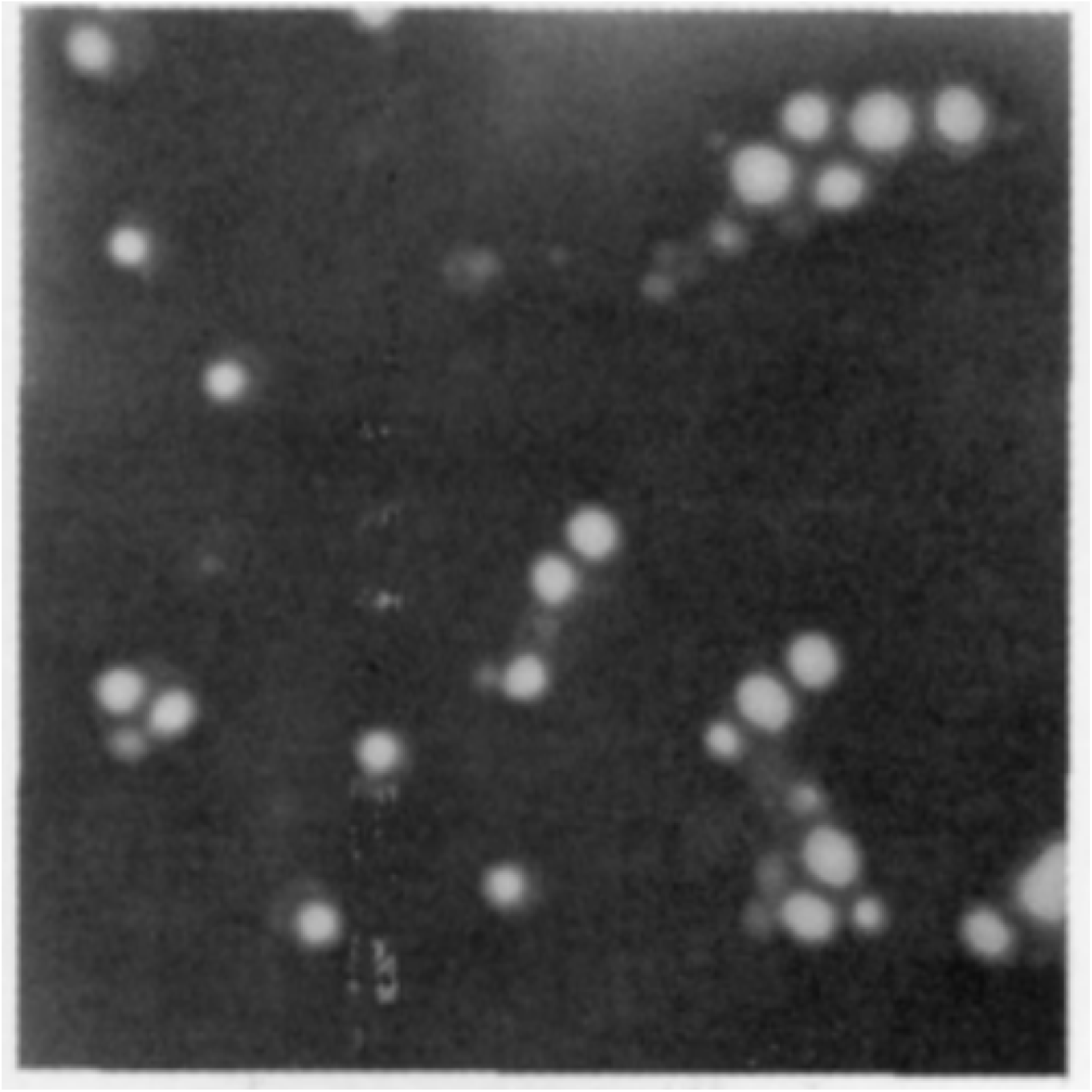
TEM analysis.

The preparation methods of the nanoparticles include an emulsion solvent evaporation method [10], a nano precipitation method [11], a high pressure homogenization method [12], and the like. In this experiment, 5-Fu nanoparticles were prepared by a simple and safe nano-precipitation method. The organic solvent used in this method is low in toxicity and easy to remove, and avoids the use of a highly toxic organic solvent such as methylene chloride. The particle prepared by this method has a small particle size and good dispersibility and reproducibility.

**Figure 4.**
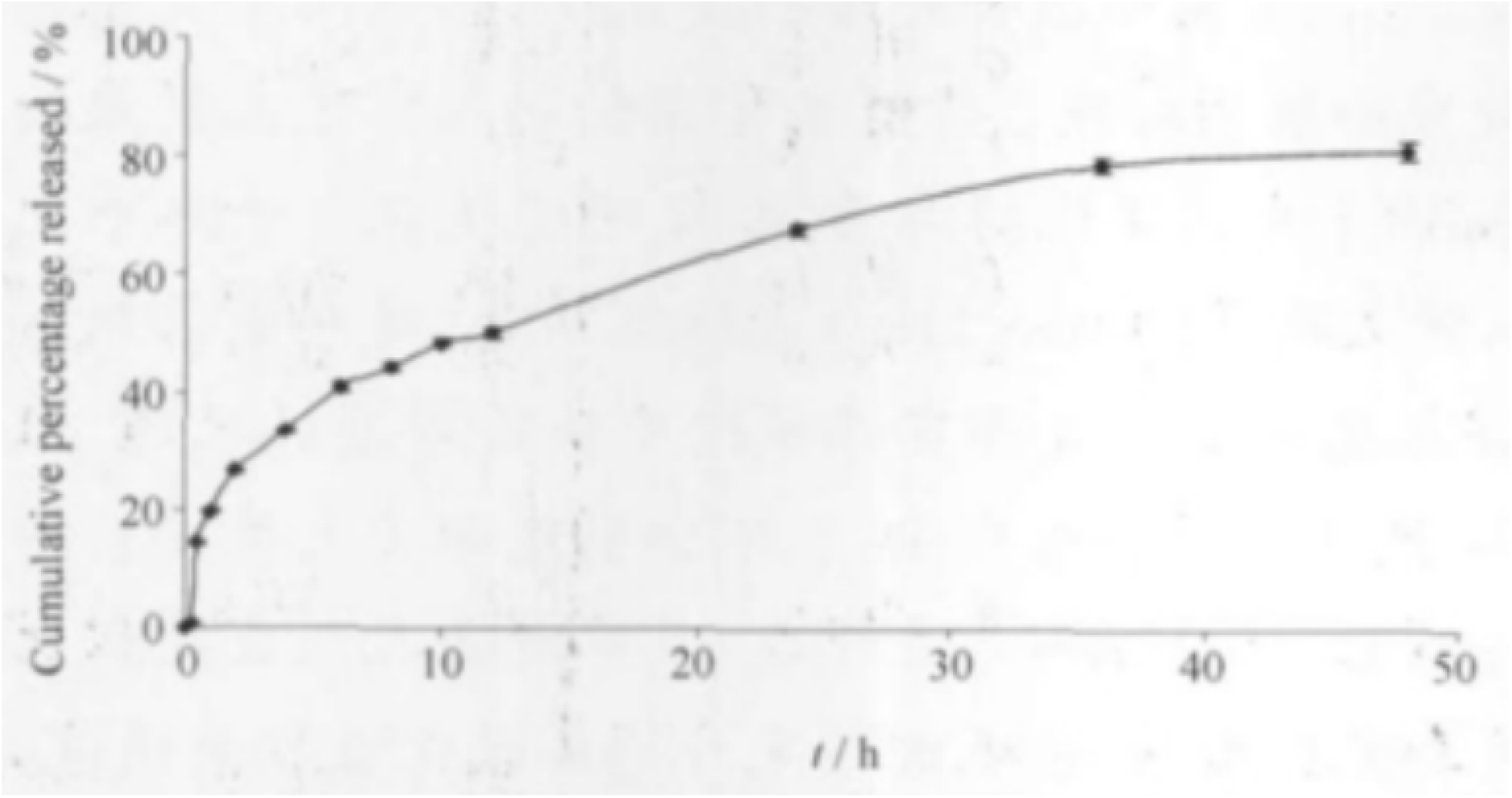
Releasing kinetics.

In this experiment, nanoparticles with a particle size of about 50 nm have been fabricated, but the release curve of the release curve is severe. At 3 h, the cumulative drug release reached 80%. The release profile of nanoparticles with a particle size of about 120 nm shows that nanoparticles with a particle size of about 120 nm have a cumulative drug content of more than 80% at 48 h, and have good sustained-release properties. It can be seen that the size of the particle size affects the release of the drug from the nanoparticles. Nanoparticles with larger particle size have a longer drug release time and can achieve a good sustained release effect^1-3^.

When determining the drug encapsulation rate, it is necessary to separate the free drug from the encapsulated drug. Common methods include dialysis, gel column chromatography, and ultrafiltration. The dialysis method takes a long time, the gel column chromatography equipment is complicated, the ultrafiltration method is expensive, and the low temperature ultracentrifugation method has simple operation and high separation ^4-15^free drugs and nanoparticles. The recovery experiment results show that the method is reliable.

Investigating the factors affecting the formulation process, the organic phase composition, temperature, water phase concentration, ratio of drug amount to polymer amount, and organic phase addition method have an effect on the encapsulation efficiency of 5-FPN. The first four factors are more affected. Significant. In this experiment, the results of the validation experiment showed that the reproducibility of the optimal prescription was good. 5-Fu is slightly soluble in water, which may be the reason for the low encapsulation efficiency. It can be seen from the experiment that the stirring time of the particles and the excessive evaporation time will reduce the encapsulation efficiency of the particles. Therefore, this experiment intends to further prepare the nanoparticles into freeze-dried powders by using the freeze drying technology to improve the stability of the particles^1-6, 16-22^.

## Conclusion

The nanoprecipitation method is simple to operate. The prepared fluorouracil PLGA-mPEG nanoparticles have small particle size and good drug release in vitro. Future efforts will focus on translating this promising technology to the clinical settings.

